# Associations of Cumulative Paternal and Maternal Childhood Maltreament Exposure with Neonate Brain Anatomy

**DOI:** 10.1101/2022.10.16.512276

**Authors:** Jetro J. Tuulari, Eeva-Leena Kataja, Linnea Karlsson, Hasse Karlsson

## Abstract

**Background:** Childhood maltreatment exposure (CME) can lead to adverse long-term consequences for the exposed individual. Emerging evidence suggests that the long-term effect of CME may be transmitted across generations, starting already during prenatal development.

**Methods:** In this study, we measured brain grey and white matter volumes from MR images in 62 healthy neonates at 2–5 weeks of gestation corrected age and obtained Trauma and Distress Scale (TADS) questionnaire data from both parents.

**Results:** We found that paternal CME associated positively with neonate supratentorial grey matter volumes while the association for the maternal TADS scores was not statistically significant. Maternal pre-pregnancy BMI associated with supratentorial white matter volumes, but not with parental CME.

**Conclusions:** We are the first to report that paternal CME is linked with variation in newborn cortical volume. Our results imply an intergenerational transmission of paternal CME to offspring. Elucidating the later relevance of these associations and mechanisms involved remains an enticing avenue for future studies.

## Introduction

Childhood trauma via childhood maltreatment exposure (CME) of varying degrees is common and represents a formidable risk factor for later psychiatric health, increasing risk for example to personality, mood, and substance use disorders and even psychosis ^1^. Emerging results have provided valuable insights on how CME exerts physiological, psychological and social traits that can not only increase health risks to the exposed individual, but also convey effects to the offspring of the exposed individual via intergenerational transmission of traits that affect the physiology and brain development of CM-exposed individuals’ offspring ^2^.

A recent study by Moog et al. put forward a hypothesis that the process of intergenerational transmission may start already during intrauterine life, and that the developing fetal brain may represent a target of particular interest ^3^. They found that maternal CME was associated with lower child intracranial volume, and cortical gray matter, and were able to demonstrate this effect in group comparison (high vs. low maternal CME) and in linear regression model with the severity of the CME using the childhood trauma questionnaire (CTQ). The potential mechanisms of the mother-to-child transmission of CME includes direct effects via altered gestational biology (epigenetic, endocrine, immune / inflammatory mechanisms) and indirect effects that occur postnatally ^4^.

We recently reported that paternal cumulative early life stress positively associates with infant offspring’s white matter integrity of the brain as measured by fractional anisotropy ^5^, which implies more advanced brain development with increasing paternal CME. This is an intriguing finding as after conception the father does not have a direct physiological connection with the developing fetus, and the paternal effects on the offspring during pregnancy are limited to indirect influences via the mother. One potential mechanism, although not confirmed in humans, could be the transmission of epigenetic marks in paternal germ line (sperm cells) ^6–8^. Neuroimaging close to birth during early neonatal period minimize the role of postnatal exposures, and indeed, this approach has been used by many investigators in the past to study prenatal exposures via the mother ^4,9^. Importantly, neonatal magnetic resonance imaging (MRI) is well suited for addressing intergenerational effects of CME of both parents.

The current study was conducted in a prospective cohort, consisting of 62 mother-father-child trios assessed prospectively from early gestation through birth until early neonatal period. This setting allowed us to conceptually replicate the analyses from the prior study of maternal CME ^3^ and to extend the approach to paternal CME that we used to predict neonatal grey and white matter volumes. Here, we used the total brain volumes as a proxy for the achieved neuronal growth during in utero development. Correspondingly, we hypothesized that maternal CME would have negative associations to brain volumes and that paternal associations would be positive in line with our prior work implicating “more advanced” brain matter maturation with increasing levels of paternal CME ^5^.

## Methods

The study was conducted in accordance with the Declaration of Helsinki and was approved by the Ethics Committee of the Hospital District of Southwest Finland (15.3.2011 §95, ETMK: 31/180/2011). We followed the Strengthening the Reporting of Observational Studies in Epidemiology (<u>STROBE</u>) reporting guideline.

**Participants** were recruited at gestational week (gwk) 12 from maternity clinics in Southwest Finland from 2011 to 2015 to take part in the FinnBrain Birth Cohort Study (http://www.finnbrain.fi), which was established to prospectively investigate the effects of early life stress (ELS), including prenatal stress exposure, on child brain development and health ^10^. The cohort entails 3808 families and included full trios (mother, father, child) on approximately half of the families.

**The demographics and questionnaire data** were obtained over the prenatal period and, although not used here, continued postnatally. Information about the parents’ CME was collected using the Trauma and Distress Scale (TADS) at gwk 12 ^11^. The TADS comprises of five core domains: emotional neglect, emotional abuse, physical neglect, physical abuse, and sexual abuse. In this study, we calculated the cumulative exposure to ELS events of the infants’ fathers and mothers by the age of 18 years (sum over the individual factor scores). Maternal depressive symptoms during pregnancy were measured using the Edinburgh

Postnatal Depressive Scale (EPDS) at gwk 24 ^12^. Maternal measures of pre-pregnancy body mass index (BMI) and level of education (low: up to 12 years of formal education; middle: 12-15 years; and high: >15 years) were available from questionnaires and the Finnish Medical Birth Register held by the National Institute for Health and Welfare. Other key parental demographics were obtained by questionnaires ^10^ and birth related factors from the Finnish Medical Birth Register (Table 1).

**Table 1.** Demographics of study participants.

| Table 1. Demographics of study participants |  |  |  |  |  |
| --- | --- | --- | --- | --- | --- |
|  |  | Mean | SD | Min | Max |
| Infant characteristics |  |  |  |  |  |
| Gestational age at birth (weeks) |  | 39.9 | 1.1 | 37.9 | 42.1 |
| Age from birth (days) |  | 26.4 | 6.6 | 11 | 43 |
| Conceptional age at scan (days) |  | 305 | 7.7 | 291 | 323 |
| Infant sex |  | 31 boys |  |  |  |
|  |  | 31 girls |  |  |  |
| Birth weight (grams) |  | 3461 | 438 | 2530 | 4700 |
| Birth length (cm) |  | 50.5 | 1.9 | 44 | 56 |
| Head circumference at birth (cm) |  | 35.0 | 1.3 | 32.5 | 37.5 |
| Parental characteristics |  |  |  |  |  |
| Maternal age at birth (years) |  | 29.9 | 4.6 | 19.0 | 41.0 |
| Paternal age at birth (years) |  | 31.2 | 4.8 | 20.0 | 43.0 |
| Maternal education level * | low | 18 |  |  |  |
| *1 missing* | mid | 18 |  |  |  |
|  | high | 25 |  |  |  |
| Paternal education level * | low | 29 |  |  |  |
| *1 missing* | mid | 15 |  |  |  |
|  | high | 18 |  |  |  |
| Maternal EPDS score at gwk 24 |  | 4.93 | 4.81 | 0.00 | 25.00 |
| Maternal pre-pregnancy BMI |  | 24.48 | 3.81 | 17.48 | 34.42 |
| Maternal TADS sum score |  | 8.2 | 8.5 | 0 | 49 |
| Paternal TADS sum score |  | 8.4 | 7.8 | 0 | 38 |
| Brain volumes |  |  |  |  |  |
| Supratentorial grey matter (mm<sup>3</sup>) |  | 156 831 | 18 119 | 115 643 | 192 161 |
| Supratentorial white matter (mm<sup>3</sup>) |  | 168 030 | 13 672 | 134 787 | 198 148 |
| **Footnote** |  |  |  |  |  |
| **\*Educational levels: Low&Mid = Upper secondary school, vocational school or lower & University of applied sciences, High = University** |  |  |  |  |  |

**Neuroimaging visits** were performed for 180 neonates at gestation-corrected age 2-5 weeks ^13,14^. The families were recruited via telephone. Exclusion criteria for infants were occurrence of any perinatal complications with potential neurological consequences, less than 5 points in the 5-minute Apgar score, previously diagnosed central nervous system anomaly or prior clinical magnetic resonance scan at peripartum due to clinical indications, gestational age of less than 32 weeks, or birth weight less than 1500 g. These criteria were confirmed through a structured phone interview. Families were provided oral and written information about the study, and the parents provided written consent to participate on behalf of their child.

**MRI acquisition protocol** is provided in our previous publications ^13,14^. Participants were scanned with a Siemens Magnetom Verio 3 T scanner (Siemens Medical Solutions, Erlangen, Germany) during natural sleep. The 40-60-min imaging protocol included an axial PD-T2-TSE (Dual-Echo Turbo Spin Echo) sequence (repetition time [TR]: 12 070 ms, effective echo times [TE]: 13 ms and 102 ms) and a sagittal 3D-T1 MPRAGE (Magnetization Prepared Rapid Acquisition Gradient Echo) sequence (TR: 1900 ms, TE: 3.26, inversion time: 900 ms) with whole brain coverage and isotropic voxels of 1.0 mm^3^ for both sequences. All brain images were assessed for incidental findings by a pediatric neuroradiologist ^15^. We were able to acquire structural MRI data from 126 of 180 infant participants.

**Preprocessing of the structural MRI data** was performed via two pipelines. We used infant FreeSurfer to segment the cerebral cortex, supratentorial white matter and ventricle volumes. 102 / 126 of the data had good segmentation results. For the subcortex, we used the output from our optimized infant pipeline ^16^.

**The final sample for the current study, included** participants that had successfully processed neonatal neuroimaging data, paternal and maternal data (including their TADS scores and relevant demographics) were available for 62 trios (fathers, mothers, and infants). The demographics are described in Table 1. There was missing information (with 1 to 6 values missing; see Table 1 footnotes) in the following maternal variables: level of education, TADS score, EPDS, prepregnancy BMI, use of SSRIs and SNRIs, alcohol use, and mode of delivery. Missing information was imputed using mean (continuous variables) or mode (categorical variables) values of the variables. The flowchart of how the final sample was reached is shown in Figure 1.

**Figure 1.**
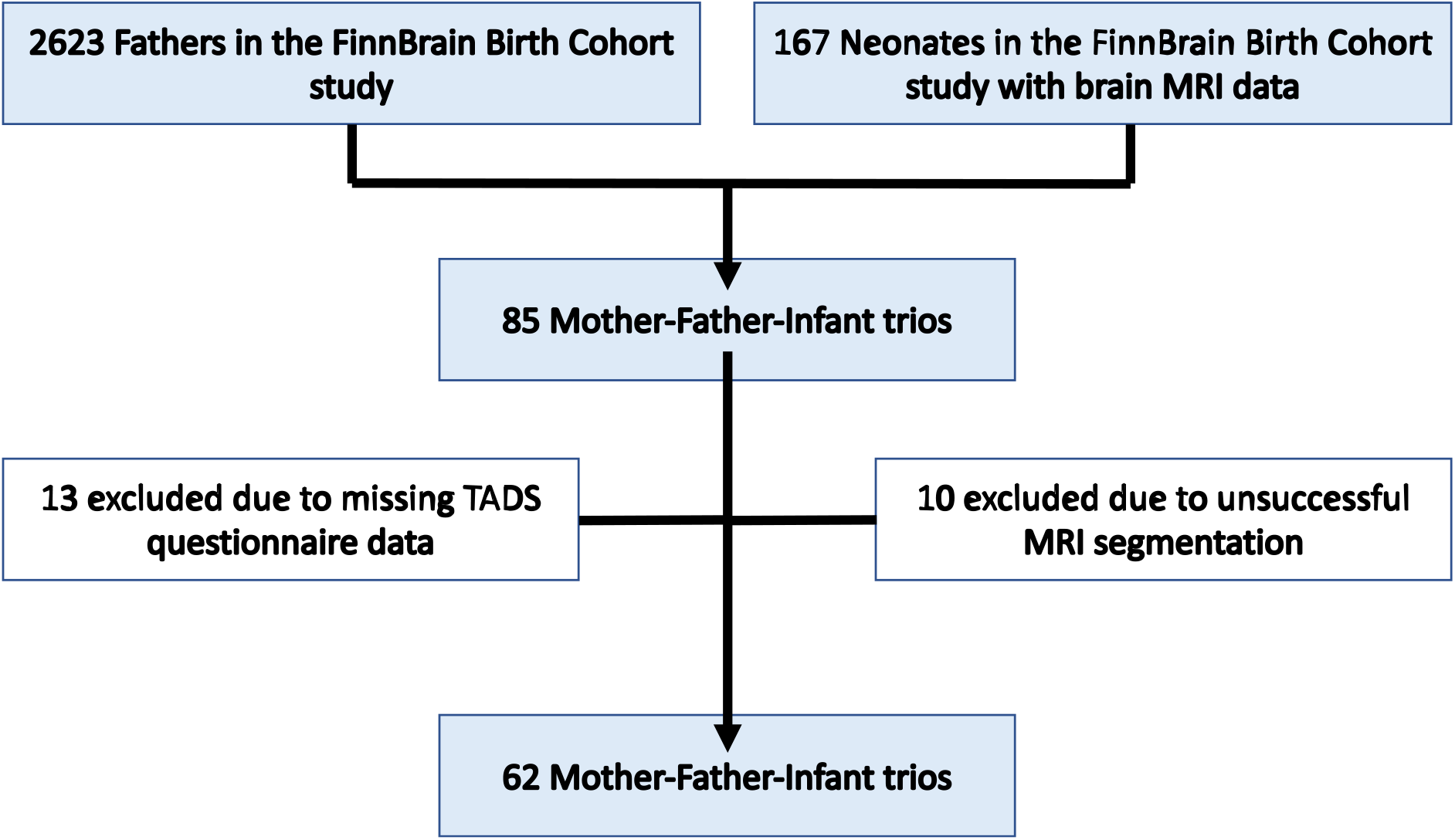
Flowchart of the Study enrolment.

Risk of bias assessment is as follows: the exposed and non-exposed parents were drawn from the same population (low risk of bias), TADS questionnaire probes CME retrospectively in a single time point (intermediate risk of bias), the outcome of interest, neonatal brain volumes, were not present at the start of the study or at the time of exposure (low risk of bias), the statistical models controlled for important confounders (low risk of bias).

**Potential maternal confounding factors** were also assessed. Seven mothers reported drinking 1 to 2 drinks more seldom than monthly. Three mothers reported smoking 5 to 10 cigarettes per day; other smokers reported quitting when they became pregnant. There was no reported illicit drug use in the current sample. Four mothers reported using SSRI / SNRI medication during pregnancy. Additionally, obstetric risk was formalized in a binary value indicating the presence of one or more of the following risk factors extracted from the medical record: hypertension, diabetes, severe anemia, severe infection, or vaginal bleeding (N = 11/62 had obstetric risk).

### Statistical Analysis

Supratentorial grey matter (GM) and white matter (WM) were the main outcome variables in the current study since they are the components of the nervous system that most clearly relate to brain growth during *in utero* brain development. For completeness, we also report result with subcortical brain volumes, total brain volume (cortical + subcortical + white matter volumes), ventricular volume, and intracranial volume in the supplementary materials.

We used R version 4.1.2 (2021-11-01) for performing all statistical analyses. The analyses relied on the following libraries: library(‘car’) #Partial plot, library(‘tidyverse’) #Data handling, library(‘rsq’) #R-squared, library(‘lm.beta’) #to get standardised regression coefficients, library(‘readr’) # to import data, library(‘olsrr’) # Residual testing, library(‘jtools’) # regression QC, library(‘sandwich’) # regression QC, library(‘apaTables’) # for exporting the results, library(‘psych’) # for getting descriptive statistics. We had interest to both maternal and paternal CME and multiple factors are known to be relevant for neonatal brain volumes. For conciseness we used a single regression model as our primary model:

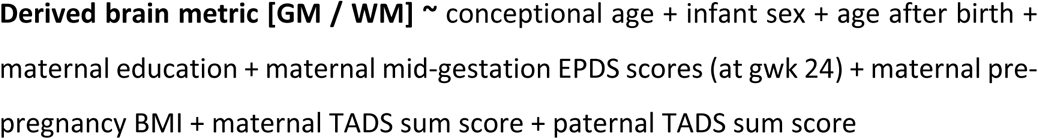

The linear regression models were checked for homoscedasticity, normality of residuals (residual plots, Q-Q plot), and collinearity with variation inflation factor (VIF). We report raw p values throughout the manuscript, but considered Bonferroni corrected p values of p < 0.025 (corrected over two regression models using supratentorial GM and WM) as statistically significant. Finally, we performed a sensitivity analysis where we additionally included binary variables of maternal smoking, SSRI/SNRI medication, alcohol use and obstetric risk.

## Results

We found that paternal CME associated positively with neonate total cortical and subcortical grey matter volumes while the negative association for the maternal trauma scores were not significant (p = 0.37) (Figure 2, Table 2). Conceptional age (positive association) and neonate sex (larger volume in boys) were expectedly predictors of total grey matter volume while the associations were not significant for other predictors. It is noteworthy that beta estimates measures for paternal TADS scores were slightly more than half of those for gestational age (in weeks). According to regression model beta estimates, this corresponds to 0.5 ml of additional grey matter volume for each TADS score. The results remained similar in sensitivity analyses (main regression model for effects of paternal TADS association beta 588.3, SE 243.0, p = 0.019; sensitivity analysis beta 647.21, SE 255.88, p = 0.0149), and none of the added variables had significant main effects.

**Table 2.**
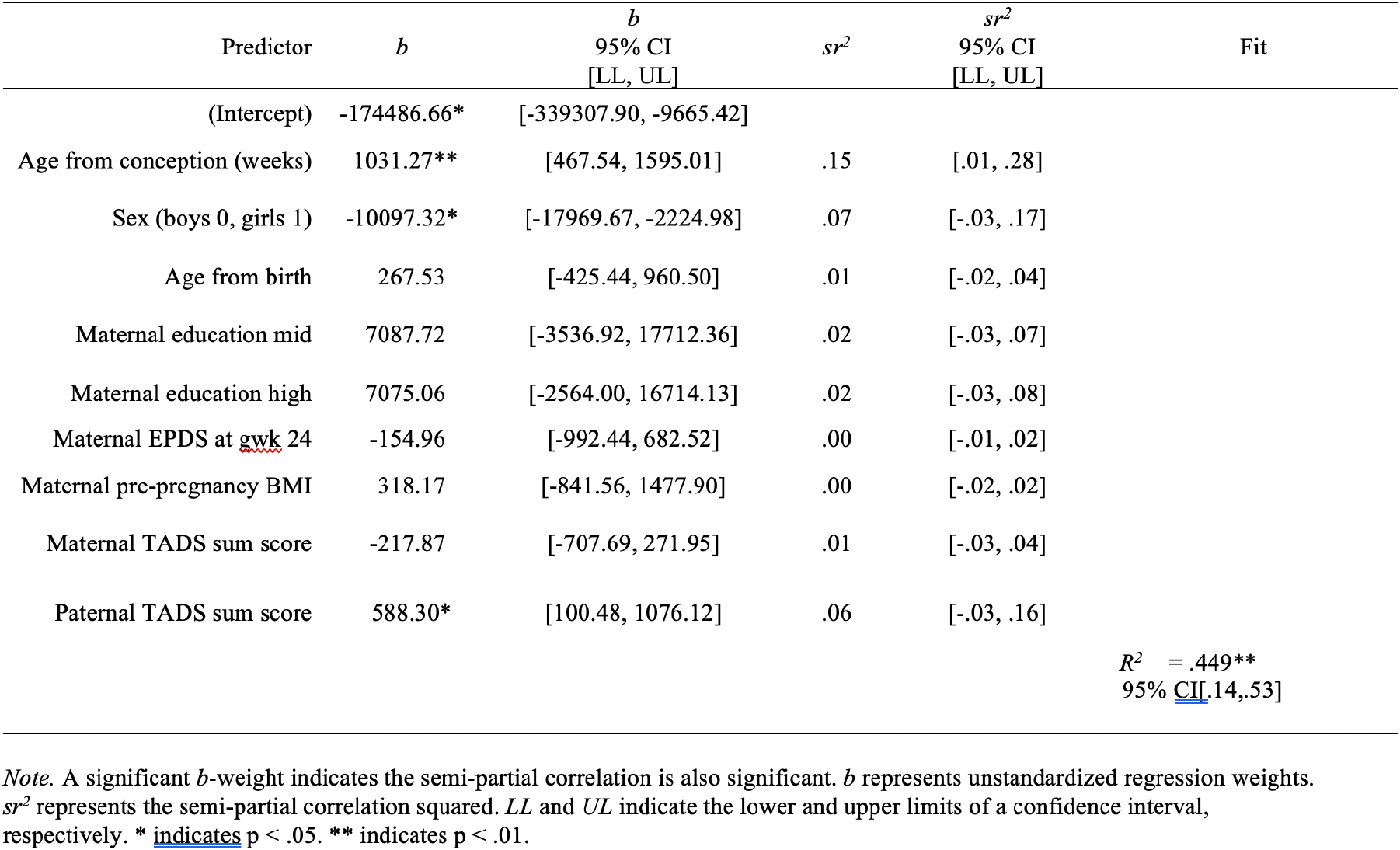
Regression model testing the association between parental TADS scores and neonatal cortical grey matter volume. All VIFs < 1.38, Adjusted R-squared: 0.3521. Created with R library (‘apaTables’).

**Figure 2.**
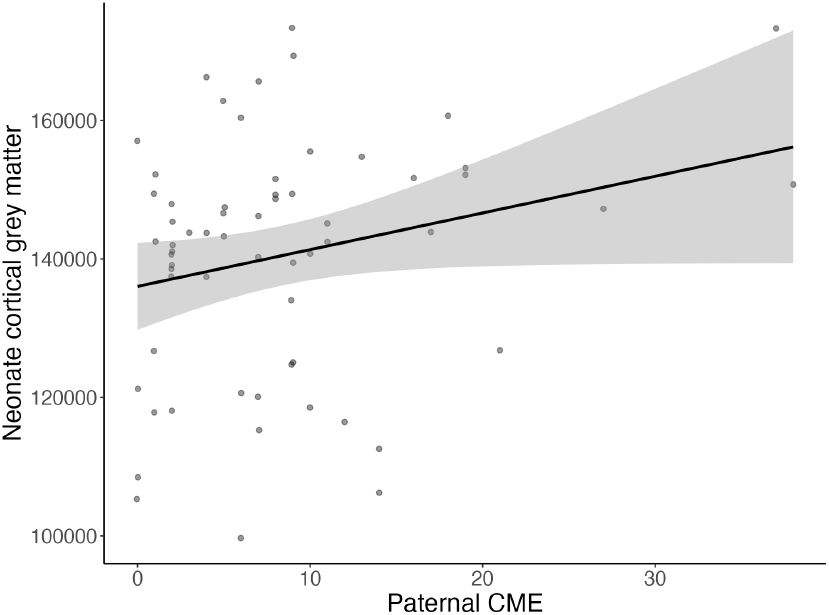
Scatter plot showing the association between neonate cortical grey matter volumes and paternal childhood maltreatment exposure (CME). Zero order Spearman correlation rho = 0.23, p = 0.077; partial Spearman correlation rho = 0.37, p = 0.006 (conditioned on infant sex, infant age at conception, and maternal education, pre-pregnancy BMI, CME scores and EPDS scores at mid gestation, in line with the main regression models, see Tables 2 and 3).

For the total supratentorial white matter volume we did not find associations with paternal or maternal TADS scores (Table 3). Conceptional age (positive association) and neonate sex (larger volume in boys) were expectedly predictors of total grey matter volume. In addition, maternal pre-pregnancy BMI associated positively with white matter volumes while the associations were not significant for other predictors. It is noteworthy that beta estimates measures for maternal pre-pregnancy BMI were twice as high as those for gestational age (in weeks). According to regression model beta estimates, this corresponds to 1.0 ml of additional white matter volume for each increase in maternal BMI. The results remained similar in sensitivity analyses (main regression model for effects of maternal pre-pregnancy BMI beta 1135.90, SE 440.8, p = 0.013; sensitivity analysis beta 1345.20, SE 486.88, p = 0.008), and none of the added variables had significant main effects.

**Table 3.** Regression model testing the association between parental TADS scores and neonatal supratentorial white matter volume. All VIFs < 1.38, Adjusted R-squared: 0.3499. Created with R library (‘apaTables’).

| Predictor | *b* | *b*<br>95% CI<br>[LL, UL] | *sr*<sup>2</sup> | *sr*<sup>2</sup><br>95% CI<br>[LL, UL] | Fit |
| --- | --- | --- | --- | --- | --- |
| (Intercept) | -32002.94 | [-157785.35, 93779.47] |  |  |  |
| Age from conception (weeks) | 579.11** | [148.90, 1009.32] | .08 | [-.02, .18] |  |
| Sex (boys 0, girls 1) | -12636.97** | [-18644.71, -6629.24] | .19 | [.04, .35] |  |
| Age from birth (weeks) | -79.22 | [-608.05, 449.62] | .00 | [-.01, .01] |  |
| Maternal education mid | 7792.50 | [-315.63, 15900.64] | .04 | [-.03, .11] |  |
| Maternal education high | 4778.92 | [-2577.08, 12134.92] | .02 | [-.03, .07] |  |
| Maternal EPDS at gwk 24 | -152.27 | [-791.38, 486.85] | .00 | [-.02, .02] |  |
| Maternal pre-pregnancy BMI | 1135.90* | [250.86, 2020.94] | .07 | [-.03, .17] |  |
| Maternal TADS sum score | -122.42 | [-496.22, 251.38] | .00 | [-.02, .03] |  |
| Paternal TADS sum score | 233.91 | [-138.37, 606.18] | .02 | [-.03, .07] |  |
| | | | | | $R^2 = .447^{**}$<br>95% CI [.14, .52] |

*Note.* A significant *b*-weight indicates the semi-partial correlation is also significant. *b* represents unstandardized regression weights. *sr*<sup>2</sup> represents the semi-partial correlation squared. *LL* and *UL* indicate the lower and upper limits of a confidence interval, respectively.
\* indicates $p < .05$ . \*\* indicates $p < .01$ .

## Discussion

To the best of our knowledge, this is the first study to describe an association between paternal childhood maltreatment and their offspring neonate supratentorial grey matter volume. In line with prior reports ^17^, maternal CME was negatively associated with neonate total grey matter volume – although, this association did not reach statistical significance.

For interpretation of the findings a division to direct and indirect intergenerational modes of transmission is useful ^4^. During gestational CME-related physiological changes may be conveyed directly to the developing offspring via the feto-placental signaling. In a physiological sense, this is the only direct pathway of intergenerational transmission, and it continues throughout the pregnancy. In contrast, fathers prenatal influence occurs ‘directly’ only at conception, and later only indirectly via the wellbeing of the mother. Second, altered CME-related gestational biological processes may indirectly affect offspring neurodevelopment by impairing the priming of postnatal parental behavior and wellbeing, including mental health (indirect pathway of intergenerational transmission). Both parents share this route of intergenerational transmission, and crucially this influence is very short in studies using neonatal neuroimaging – as in the current study and one prior report focusing on maternal CME ^3^.

Fathers influence their offspring brain development directly only at conception. It could be speculated that these findings may be explained by direct paternal genetic transmission or paternal early life gene-environment correlation that influence brain development. Alternatively, since CME does not change DNA, the attention is drawn to epigenetics of the male germ line. Indeed, there are animal studies that implicate that intergenerational, and even transgenerational inheritance, occur through male germ cell line epigenetic modifications ^2,18,19^.The sample size of the current study is not sufficient to formally test the indirect effects of paternal influence on maternal wellbeing, and we lack the biological samples needed to test the epigenetic mechanisms. This remains an important avenue for future studies, especially those with new prospective data collections. Importantly, since the CME can be quantified retrospectively, replication studies of our results may also be conducted in existing data sets by collecting additional questionnaire data ^20^.

The effects of CME are commonly expected to be harmful also within the context of intergenerational transmission, but critically, there are no empirical studies to confirm this. Indeed, we found that paternal CME associated with larger brain volumes and our prior report with similar setting found that paternal CME associated with higher white matter integrity. Both brain metrics are commonly associated with ‘more advanced’ brain development. We could interpret this as faster early brain maturation with ultimately lower adult brain size as proposed by stress acceleration hypothesis ^21^. Alternatively, increased brain maturation may be related to differential susceptibility to environmental factors ^22,23^. Additional studies that include later postnatal outcomes are needed to assess the practical implications to child outcomes.

This study has some limitations. In line with the prior report on maternal CME ^17^, the study population was not enriched for CME or clinical groups. The sample size was relatively small, which is common in neonatal scans, but carries important limitations to generalizability and statistical power. The retrospective assessment of CME may result in recall bias and is unable to capture very early CM exposure.

## Conclusions

In conclusion, our results imply that neonatal cortical brain volume of the offspring is related to paternal early life stress exposure as measured by a sum of childhood maltreatment scores. The link of paternal CME to offspring brain development is a novel finding, and an additional factor to consider in intergenerational transmission of CME – both for future mechanistic studies and eventual development of interventions. Retrospective measures of parental CME are obtainable at any time, and this creates high hopes that future studies can replicate findings with the use of existing data sets.

## Acknowledgements

We would like to warmly thank all FinnBrain families that participated to the study. We would also like to thank the research team: Satu Lehtola for her help in data collection, Maria Lavonius for her help in recruiting the participants, Jani Saunavaara for implementing the MRI sequences, John D. Lewis for his help in planning the sequences and data preprocessing, Riitta Parkkola for reviewing the MR images for incidental findings.

## Author contributions

JJT planned the analytical approach and performed the data analyses. LK and HK planned and funded the MRI measurements, established the FinnBrain Birth Cohort and built the infrastructure for carrying out the study. All authors participated in conceptualizing the study, writing the manuscript and accepted the final version

## Funding

Jetro J. Tuulari

Suomen Aivosäätiö; Sigrid Jusélius Foundation; Emil Aaltonen Foundation; Finnish Medical Foundation; Alfred Kordelin Foundation; Juho Vainio Foundation; Turku University Foundation; Hospital District of Southwest Finland; State Grants for Clinical Research (ERVA); Orion Research Foundation, Signe and Ane Gyllenberg Foundation.

Eeva-Leena Kataja

Academy of Finland. Grant number: #346790

Signe and Ane Gyllenberg Foundation

Linnea Karlsson

Academy of Finland, Grant/Award Number: NNN

Turku University Foundation; Hospital District of Southwest Finland; State Grants for Clinical Research (ERVA);

Brain and Behavior Research Foundation;

Päivikki and Sakari Sohlberg Foundation.

Hasse Karlsson

Academy of Finland, Grant/Award Number: NNN

Turku University Foundation; Hospital District of Southwest Finland; State Grants for Clinical Research (ERVA);

